# SIRT6 activation rescues the age-related decline in DNA damage repair in primary human chondrocytes

**DOI:** 10.1101/2023.02.27.530205

**Authors:** Michaela E. Copp, Jacqueline Shine, Hannon L. Brown, Kirti R. Nimmala, Susan Chubinskaya, John A. Collins, Richard F. Loeser, Brian O. Diekman

**Affiliations:** Thurston Arthritis Research Center, University of North Carolina at Chapel Hill, Chapel Hill, NC; Joint Department of Biomedical Engineering, University of North Carolina and North Carolina State University, Raleigh, NC; Department of Pediatrics, Rush University Medical Center, Chicago, IL; Department of Orthopedic Surgery, Thomas Jefferson University; Division of Rheumatology, Allergy, and Immunology, University of North Carolina

**Keywords:** DNA Damage, Sirtuins, MDL-800, DNA Repair, Cartilage, Osteoarthritis, Aging, Comet assay

## Abstract

While advanced age has long been recognized as the greatest risk factor for osteoarthritis (OA), the biological mechanisms behind this connection remain unclear. Previous work has demonstrated that chondrocytes from older cadaveric donors have elevated levels of DNA damage as compared to chondrocytes from younger donors. The purpose of this study was to determine whether a decline in DNA repair efficiency is one explanation for the accumulation of DNA damage with age, and to quantify the improvement in repair with activation of Sirtuin 6 (SIRT6). Using an acute irradiation model to bring the baseline level of all donors to the same starting point, this study demonstrates a decline in repair efficiency during aging when comparing chondrocytes from young (≤45 years old), middle-aged (50-65 years old), or older (>70 years old) cadaveric donors with no known history of OA or macroscopic cartilage degradation at isolation. Activation of SIRT6 in middle-aged chondrocytes with MDL-800 (20 μM) improved the repair efficiency, while inhibition with EX-527 (10 μM) inhibited the rate of repair and the increased the percentage of cells that retained high levels of damage. Treating chondrocytes from older donors with MDL-800 for 48 hours significantly reduced the amount of DNA damage, despite this damage having accumulated over decades. Lastly, chondrocytes isolated from the proximal femurs of mice between 4 months and 22 months of age revealed both an increase in DNA damage with aging, and a decrease in DNA damage following MDL-800 treatment.

## Introduction

Osteoarthritis (OA) is a chronic joint disease affecting approximately 13% of the US adult population and is characterized by the degradation of articular cartilage, synovial inflammation, and subchondral bone remodeling^1–3^. As no disease-modifying therapies for OA have been FDA approved to date^4^, the main options available to OA patients are pain management and eventual total joint replacement, leading to extensive societal and economic burdens^5^. While a number of risk factors have been associated with OA – obesity, biological sex, joint injury, and genetics – the leading risk factor is older age^6^. While progress continues to be made, the biological mechanisms linking aging and OA prevalence remain largely unknown^7^.

Hypo-replicative cell types such as neurons, hematopoietic stem cells, and chondrocytes tend to accumulate sites of persistent DNA damage during aging, due at least in part to the lack of access to repair mechanisms that are only present in S phase^8–10^. As measured by the alkaline comet assay^11,12^, we showed that chondrocytes isolated from older cadaveric donors, despite no known history of OA or macroscopic cartilage damage, harbor high levels of DNA damage^13^. One objective of this study was to determine whether a reduced efficiency of DNA damage repair with aging is one potential cause of DNA damage accumulation.

Sirtuin 6 (SIRT6) is nuclear-localized NAD(+)-dependent deacetylase that has been shown to play numerous important roles in cellular processes that become dysregulated with aging^14–17^. SIRT6 quickly localizes to sites of DNA damage and initiates chromatin remodeling to facilitate the recruitment and activity of proteins involved in DNA repair^18–22^. Prior work has indicated that SIRT6 is a critical factor in joint tissue homeostasis^23–26^, and the level and/or enzymatic activity of SIRT6 decreases with age and in the context of OA^25,27^. Small molecules can be used to either increase or decrease the deacetylase activity of SIRT6: MDL-800 is an allosteric activator that increases activity by ~22-fold^28^, whereas EX-527 is an inhibitor that stabilizes the closed conformation of sirtuins^29^ and blocks 67% of recombinant SIRT6 activity within 15 minutes^25^. The second objective of this study was to examine how modulating SIRT6 activity impacts the repair of DNA.

Prior work completed in our lab has demonstrated that primary human chondrocytes accumulate damage in a linear manner with age, predominantly driven by strand breaks to the DNA^13^. The third objective of this study was to determine the extent to which MDL-800 can reduce DNA damage that has accumulated over decades in chondrocytes from older donors. Similarly, we investigated whether murine chondrocytes show a similar increase in DNA damage with age and whether MDL-800 treatment is sufficient to reverse damage in this important model species.

In this study, we use irradiation as an acute model of DNA damage to show that the DNA repair efficiency of chondrocytes deteriorates throughout life but can be enhanced by activating SIRT6. Further, SIRT6 activation was sufficient to reduce the accumulated DNA damage that arises with aging in human and murine chondrocytes. These findings support DNA damage repair as one beneficial aspect of SIRT6 and promote the activation of SIRT6 as a possible point of intervention to mitigate age-related OA.

## Results

### Decreased DNA damage repair efficiency with aging in primary human chondrocytes

To investigate how aging impacts the repair capacity of chondrocytes, we used irradiation to apply an acute bolus of damage to cells and monitored DNA damage by the comet assay at time points of 15, 30, 45, 60, 120, and 240 minutes after damage. This irradiation model allowed us to apply nearly instantaneous damage to the cells and conduct a precise time-course study of repair by transferring the slides directly to the lysis buffer (experimental approach in Fig. 4). Importantly, chondrocytes from distinct age ranges of young (≤45 years old), middle (50-65 years old), and older (>70 years old) had a similar amount of DNA damage immediately after irradiation, indicating that this bolus of damage was sufficient to overcome the background differences in accumulated damage. The ability of the chondrocytes to resolve DNA damage from this equal starting point over the course of 4 hours was impaired in the middle-aged and older donors as compared to the young donors (Fig. 1A). The older donors had a significantly higher percentage of DNA in comet tails as compared to the middle and/or younger donors at 60, 120, and 240 minutes (p<0.05, multiple comparisons test). By 4 hours post-irradiation, most of the damage was resolved in chondrocytes from younger donors, whereas the average percentage of DNA in comet tails remained elevated for both middle-aged and older donors. Additional insight can be gained by assessing the distribution of damage within individual cells for each donor, as shown for representative young, middle, and older donors (Fig. 1B). Of note, there was a bifurcated response in the older donors, with a significant fraction of cells retaining very high levels of damage (dotted line at 60% of damage in comet tails). When quantified across all donors, 27.6% of chondrocytes in the older group retained this high level of damage at 4 hours, whereas this percentage was 12.5% and 2.6% for middle and younger donors, respectively (Fig. 1C). Analysis of chondrocytes with <15% DNA in comet tails at 4 hours showed that 68.7%, 49.6%, and 41.3% of cells from young, middle, and older donors, respectively, repaired the damage from irradiation to near-baseline levels (Fig. 5).

**Figure 1:**
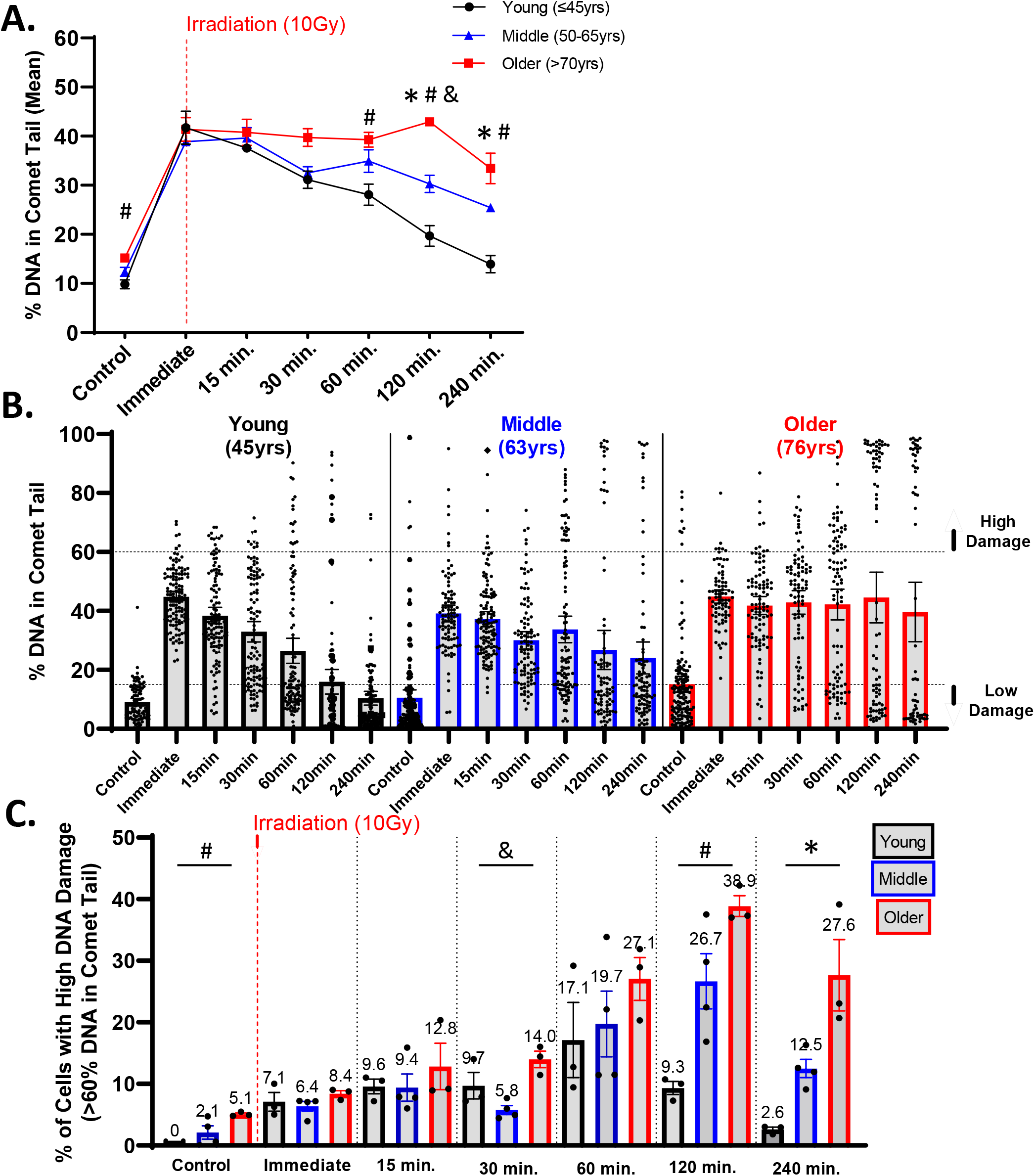
Repair after acute DNA damage in chondrocytes from different aged donors. Primary human chondrocytes from young (n=3, ≤45 years), middle-aged (n=4, 50-65 years), and older (n=3, >70 years) were prepared in gels on microscope slides and irradiated with 10 Gy (or not for control). Slides were transferred to media for repair after irradiation and then lysis buffer at various time points. (A) The percentage of DNA in comet tails for all cells were averaged for each donor, and the mean of all donors per age group is shown (mean + SEM). Repair time, age, and their interaction were significant sources of variation (2-way repeated measures ANOVA). Significant differences between groups at each time point (Tukey’s multiple comparisons test, p<0.05) are denoted by symbols: (*) = young vs. middle, (#) = young vs. old, (&) = middle vs. old. (B) Plots showing all individual cells of representative young, middle, and older donors. (C) The percentage of cells with high levels of DNA damage (>60% of DNA in comet tails). Statistics same as in B.

### SIRT6 activation and inhibition affects the repair efficiency of chondrocytes

As SIRT6 is critical in the DNA repair process of cells, we sought to study how modulating SIRT6 activity impacts the repair of DNA. Using the same irradiation and comet assay system to study repair efficiency, chondrocytes from middle-aged donors were pre-treated for 2 hours with MDL-800 (activator), EX-527 (inhibitor), or DMSO (vehicle control). Following irradiation, the slides were placed back into media baths with their respective treatments for recovery (15 to 240 minutes post-irradiation), such that the cells were receiving SIRT6 activation/inhibition for the entirety of the study (experimental approach in Fig. 4). When assessed by repeated measures two-way ANOVA without consideration of EX-527 treatment, MDL-800 treated groups showed lower DNA damage as compared to DMSO in middle-aged donors (Fig. 1A, main effects p-value = 0.005). Similarly, when DMSO and MDL-800 were compared in chondrocytes from older donors (>70 years), there was reduced damage with MDL-800 treatment at 30, 60, 120, and 240 minutes (Fig. 6A). Further, MDL-800 reduced the percentage of cells with high damage (>60% DNA in comet tails) at 4 hours from 20.1% to 4.9% (Fig. 6B). When EX-527 was also considered in the ANOVA for middle-aged donors, this inhibitor showed strong effects with significantly more DNA damage as compared to the DMSO and/or MDL-800 groups at baseline, 60, 120, and 240 minutes of repair (Fig. 2A). The all-cell plot shows a striking increase in the percentage of individual cells that retain high levels of DNA damage in the EX-527 group (Fig. 2B) – at 4 hours, 37.2% of cells still had greater than 60% of the DNA in comet tails. When comparing the percentage of cells with low levels of DNA damage, MDL-800 treatment significantly increased the likelihood that cells are able to restore near-baseline levels of damage at two and four hours post-irradiation (Fig. 5B).

**Figure 2.**
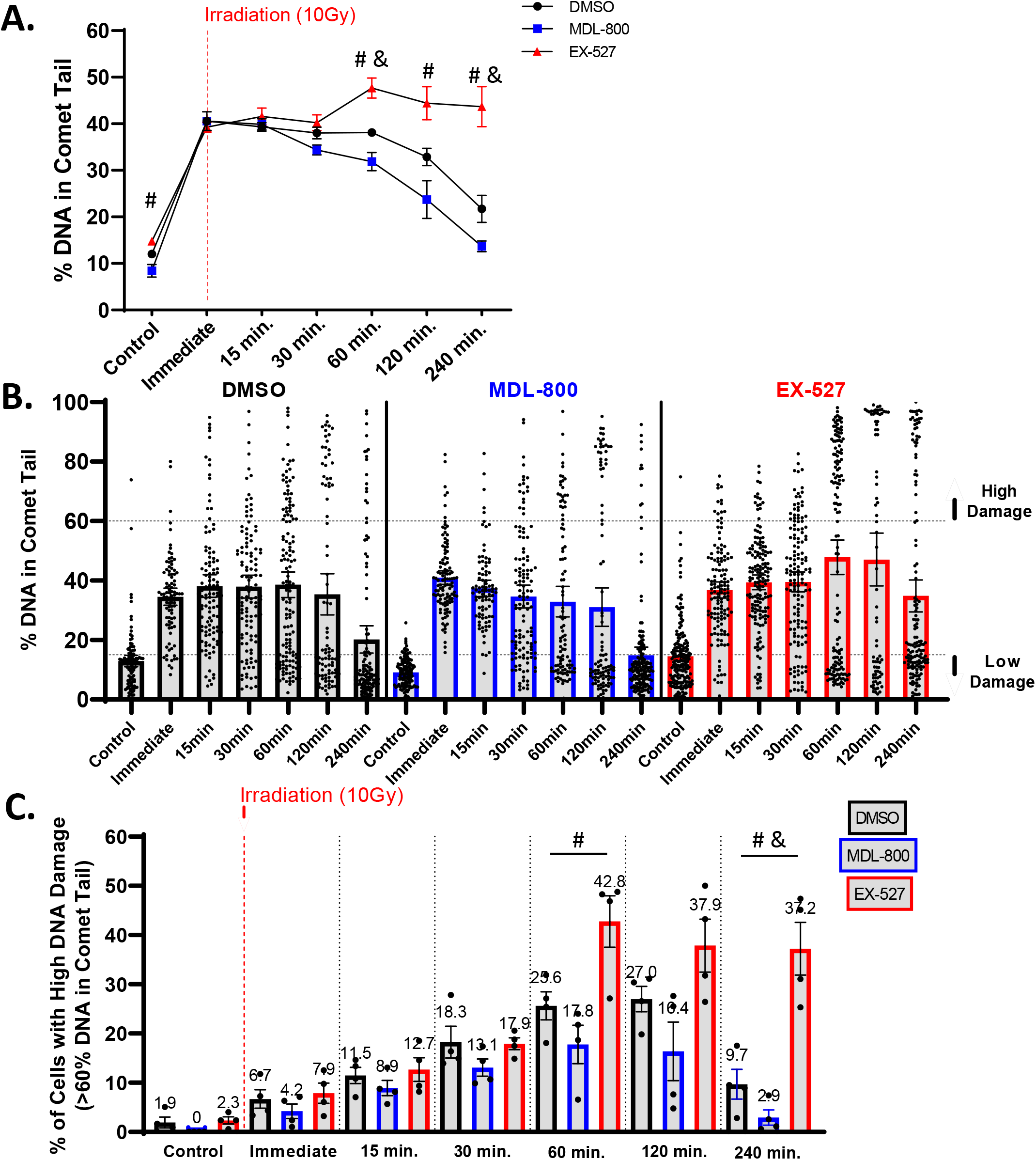
Effect of SIRT6 activation and inhibition on chondrocyte repair efficiency. Chondrocytes from middle-aged donors were pre-treated with 20 μM MDL-800, 10 μM EX-527, or vehicle (DMSO) for 2 hours before trypsinization, gel encapsulation, and irradiation. Treatment continued during the repair phase. (A) The percentage of DNA in comet tails for all cells were averaged for each donor, and the mean of all donors per condition is shown (mean + SEM). Repair time, treatment, and their interaction were significant sources of variation (2-way repeated measures ANOVA). Significant differences between groups at each time point (Tukey’s multiple comparisons test, p<0.05) are denoted by symbols: (*) = DMSO vs. MDL, (#) = MDL vs. EX, (&) = DMSO vs. EX). (B) Plots show all individual cells of a representative donor treated with DMSO, MDL-800, or EX-527. (C) The percentage of cells with high levels of DNA damage (>60% of DNA in comet tails). Statistics as in B.

### SIRT6 activation decreases DNA damage associated with older age

Having shown that SIRT6 activity affects the repair capacity of chondrocytes in response to an acute bolus of damage, we wanted to test whether MDL-800 could also repair long-standing naturally accumulated damage. In previous studies, we have established that there is higher DNA damage in chondrocytes from older donors, with a linear regression showing that donors at age 40 have ~10% DNA in comet tails and donors at age 75 have ~27%^13^. Here, we treated chondrocytes isolated from older cadaveric donors for 48 hours with either 20 μM of MDL-800 or vehicle control (DMSO). The MDL-800 treated chondrocytes showed significantly lower levels of DNA damage (mean: 11.3% of DNA in comet tails) as compared to the DMSO groups (21.3%) (Fig. 3A, p=0.0031, paired t-test).

**Figure 3:**
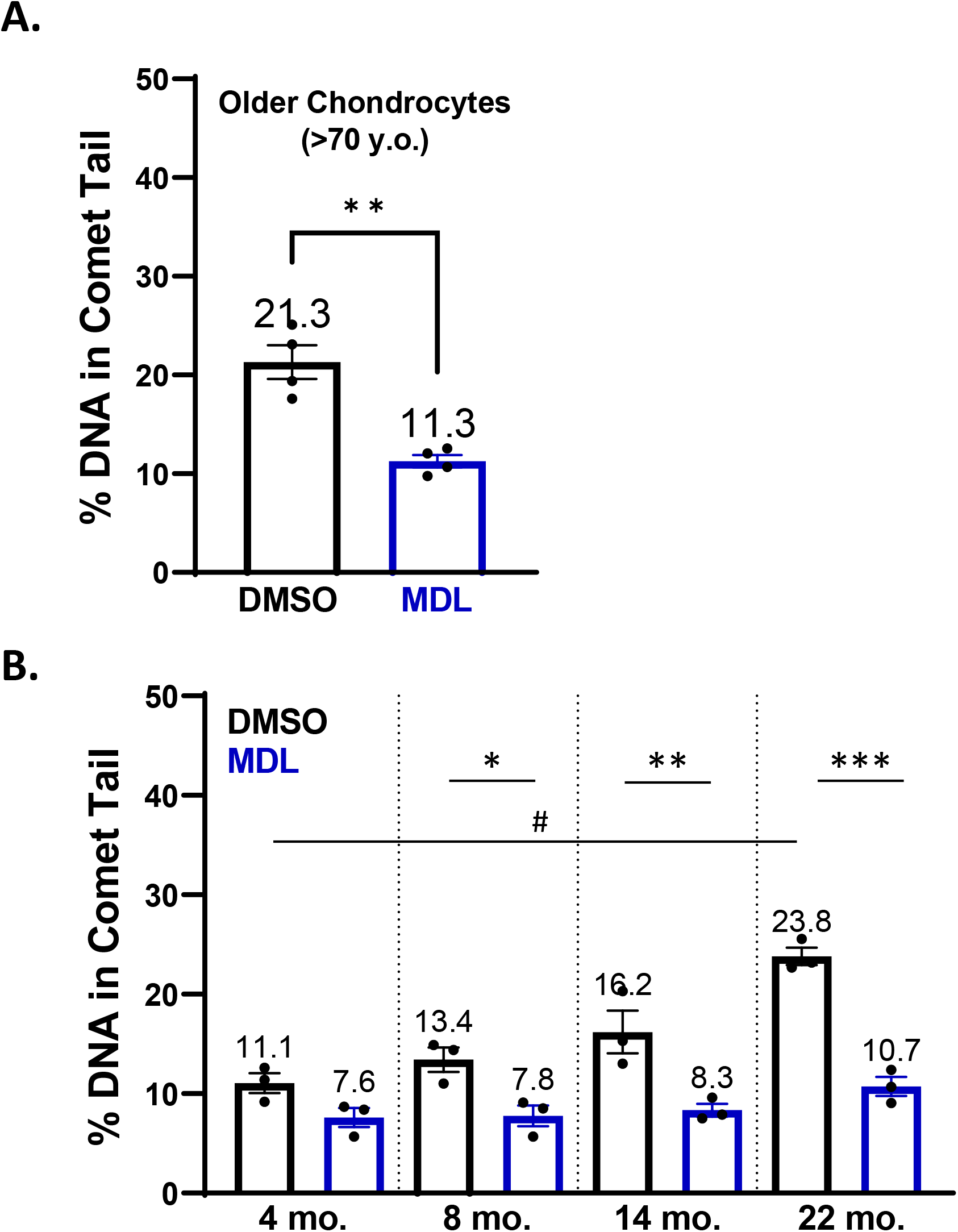
SIRT6 activation in chondrocytes from older human donors and mice. (A) Chondrocytes derived from cadaveric ankle cartilage of older donors (>70 years) were treated with 20 μM MDL-800 or vehicle (DMSO) for 48 hours. Stats by paired t-test. (B) Chondrocytes from the proximal femur of mice were isolated and treated for 48 hours with DMSO or 20 μM MDL-800. Analysis by two-way ANOVA showed significant effects of age, treatment, and their interaction. Significant differences between groups at each time point (Sidak’s multiple comparisons test, p<0.001) are denoted by # symbol, with differences between treatments denoted by *.

### MDL-800 treatment reduces DNA damage in aged murine chondrocytes

Mice are a commonly used model species for investigations of mammalian aging and thus we sought to determine whether DNA damage also accumulates with age in murine chondrocytes. Chondrocytes from the proximal femur were isolated and then treated with MDL-800 (20 μM) or DMSO control for 48 hours before comet analysis. DNA damage increased in the DMSO-treated groups with age, with the percentage of DNA in comet tails approximately doubling from 4 to 22 months of age (Fig. 3B). MDL treatment consistently lowered DNA damage in all age groups, with significant reductions at 8, 14, and 22 months of age (Fig. 3B, p<0.05, multiple comparisons test).

## Discussion

In the present study, we found that (1) advanced age negatively impacts the ability of chondrocytes to repair DNA damage, (2) modulating SIRT6 activity affects the repair capacity of chondrocytes, and (3) activating SIRT6 with MDL-800 can aid in repairing DNA damage that had accumulated over decades. The first two findings made use of irradiation to initiate a bolus of damage. This system was particularly valuable in that the level of damage immediately following irradiation was consistent across all ages and treatment groups, allowing us to directly compare the progressive reduction in DNA damage over time.

There is growing evidence that the efficiency of DNA damage repair declines with age (reviewed in ^30^ and ^31^). Previous work has largely been performed in fibroblasts and lymphocytes, but the current study confirms that aging also affects the repair of DNA damage in primary human chondrocytes. We used the alkaline comet assay to provide a sensitive and quantitative measure of DNA damage levels. Upon placement in a lysis solution, single strand breaks (SSBs), double strand breaks (DSBs), and other forms of damage (i.e., abasic sites) relax the supercoiled DNA loops of the nucleus, enabling easier movement of the DNA through the agarose gel when an electric field is applied^11,12^. As a result, damaged DNA produces a “comet tail” while intact DNA remains in the “comet head”. One advantage of this assay is the single-cell nature of the readout. This allowed us to observe that chondrocytes from older donors had a larger percentage of cells that showed very little repair and instead retained a high damage burden at four hours post-irradiation. This finding aligns with a previous study in lymphocytes that showed the primary difference with age in response to irradiation was the increased subset of cells that retained high damage^32^.

SIRT6 is involved in numerous biological processes with relevance to aging^33^, including a role in multiple DNA damage repair pathways^18,19,34–36^. Given the selectivity of MDL-800 for SIRT6^28^, we were able to show that activation of SIRT6 is sufficient to repair approximately half of the accumulated damage in chondrocytes from older human donors and from older mice. For human chondrocytes, 48 hours of treatment with MDL-800 lowered the percentage of DNA in comet tails from 21.3% to 11.3%. Based on the linear regression calculated from 25 donors ranging in age from 34 to 78 years old in Copp et al.^13^, MDL-800 treatment was able to eliminate the equivalent of ~34 years’ worth of damage is erased.

Cellular senescence is a phenotypic state characterized by stable cell cycle arrest in response to intrinsic or extrinsic stress^37^. The accumulation of senescent cells has been associated with numerous aging-related diseases and likely plays a role in OA pathogenesis^38,39^. However, less is known regarding the biological processes and environmental cues that prime chondrocytes to become senescent. Evidence supports the notion that DNA damage is a potential causative factor that drives senescence and other features of aging^40,41^, and other studies have linked DNA damage with chondrocyte dysfunction during OA^42^. Our previous work also supports a causal role for DNA damage in chondrocyte senescence, as the application of 10 Gy irradiation (which recapitulates the level of DNA damage in older donors^13^) is capable of inducing senescence in cartilage explants when paired with a mitogenic stimulus^43^. A recent study demonstrated that Sirt6 deficiency exaggerated chondrocyte senescence and OA, while intra-articular injection of adenovirus-Sirt6 or the introduction of nanoparticles releasing MDL-800 mitigated OA caused by destabilization of the medial meniscus surgery^25^. When paired with the results of the current study, an intriguing hypothesis for future work is that SIRT6 activation prevents senescence and OA through enhanced DNA damage repair capacity.

In conclusion, the findings presented here support the hypothesis that the efficiency of DNA damage repair declines with age in chondrocytes and that SIRT6 activation improves repair both in response to an acute irradiation challenge and in the context of age-related damage accumulation. These results emphasize the critical role of SIRT6 in DNA repair and support further studies investigating the use of MDL-800 (or alternative SIRT6 activators) in mitigating senescence induction and ameliorating OA development.

## Materials and Methods

### Isolation and culture of primary human chondrocytes

Primary human chondrocytes were isolated from the ankle cartilage of cadaveric donors without a history of OA and with grades between 0 and 2 on the modified Collins grade^44^. For the study presented in Fig. 1, ages of donors were in three groups: younger (40, 44, 45 years old); middle-aged (56, 56, 63 years old); older (73, 75, 76 years old). For the study presented in Fig. 2, the donors used were middle-aged (51, 54, 54, 55, 56, 56, 60, 63). For the study presented in Fig. 3A, the ages of the donors were 74, 75, 75, and 76. To isolate the primary chondrocytes, full-thickness cartilage shards were digested with 2 mg/ml Pronase (1 hour) followed by overnight incubation with 3.6 mg/ml Collagenase P at 37°C in 5% serum media^45^. The isolated chondrocytes were plated at a concentration of ~1×10^5^ cells per cm^2^ in DMEM/F12 supplemented with 10% FBS, penicillin and streptomycin, gentamicin, and amphotericin B to recover from isolation and frozen in Recovery™ Cell Culture Freezing Medium. Chondrocytes were plated for ~2-3 days before harvest for resuspension in comet gels for irradiation.

### Isolation of primary murine chondrocytes

The cartilaginous head of the proximal femur (i.e., the femoral cap of the hip) was used as the source of chondrocytes from C57BL/6 mice at various ages. Chondrocytes were isolated via pronase (2 mg/ml, 1 hour in serum-free media) and collagenase P (500 μg/ml, overnight in 10% serum) from mice aged 4, 8, 14, and 22 months of age (n=3 each). Chondrocytes were cultured for ~3 days to recover before treatment.

### SIRT6 Activation and Inhibition Treatment

The small molecule MDL-800 (Sigma) was used at a concentration of 20 μM to activate SIRT6. Conversely, EX-527 (Selleck) was used at a concentration of 10 μM to inhibit SIRT6 activity. When testing the effect of SIRT6 modulation on DNA repair following acute damage (Figure 2), primary chondrocytes were pre-treated with either DMSO (vehicle control, concentration matching the DMSO used with MDL-800), MDL-800, or EX-527 for 2 hours prior to harvest for irradiation experiments. For experiments testing the reduction of accumulated DNA damage in chondrocytes from older cadaveric donors and mice, cells were treated with DMSO or 20 μM MDL-800 for 48 hours.

### Acute Irradiation Repair Model and Comet Assay Protocol

A schematic depicting the irradiation repair model is shown in Fig. 4. After trypsinization, chondrocytes were prepared for the comet assay as described^13^, with adjustments made to measure DNA damage levels at specific time points following irradiation. Briefly, cells were mixed 1:10 with 1% low melting agarose and coated onto a Superfrost slide. The slides were placed in a media bath and irradiated with 10 Gy X-ray (RS2000 Biological Irradiator), with one slide not irradiated as a control group. The slides were moved to an incubator with their appropriate media for various amounts of time for recovery and then added to a lysis solution at the indicated time point – immediate (no recovery after IR), 15 min., 30 min., 60 min., 120 min., and 240 min. The lysis solution was prepared by mixing 2.5 M NaCl, 0.1 M disodium EDTA, 10 mM Tris base, 0.2 M NaOH, 0.1% sodium lauryl sarcosinate, and 1% Triton X-1000, and adjusting the solution to a pH of 10. After overnight incubation in the lysis solution at 4°C, the slides were added to an alkaline electrophoresis solution (200 mM NaOH, 1 mM disodium EDTA, pH > 13) for 30 minutes. Next, the slides were placed into an electrophoresis chamber and an electric field of 1 V/cm for 20 minutes was applied. Slides were washed with dH_2_O and stained with NucBlue™ (R37605; Thermo Fisher Scientific). Fluorescence images were captured with an EVOS M5000 microscope (AMF5000; Thermo Fisher Scientific). Image analysis and comet quantification were performed for approximately 100 cells per condition using the OpenComet plugin software in ImageJ.

**Figure 4:**
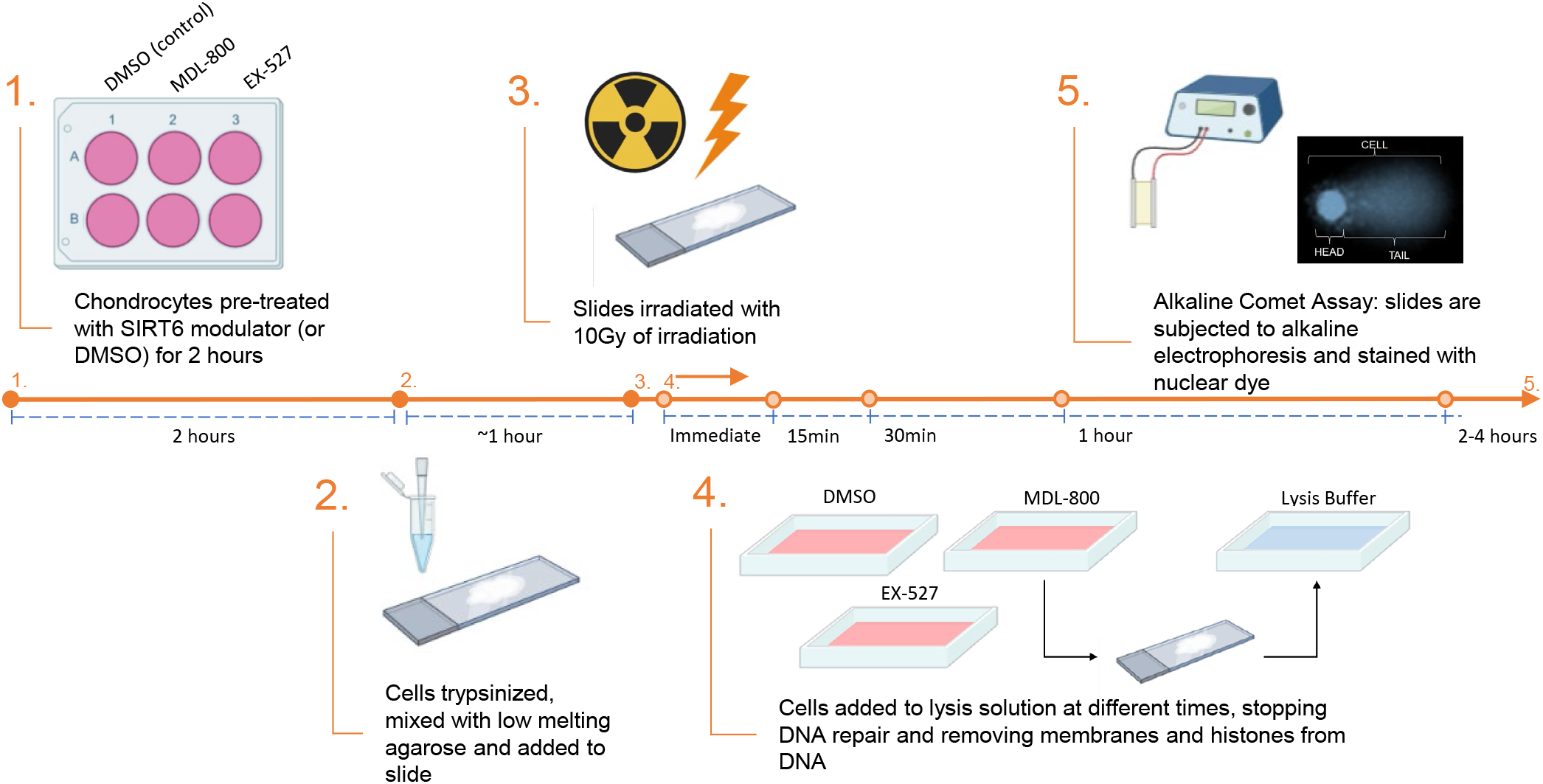
Experimental design for the results shown in Fig. 2. For the data in Fig. 1, there was no pre-treatment and steps 2-5 were completed as shown (with 10% serum media used for the repair phase).

**Figure 5:**
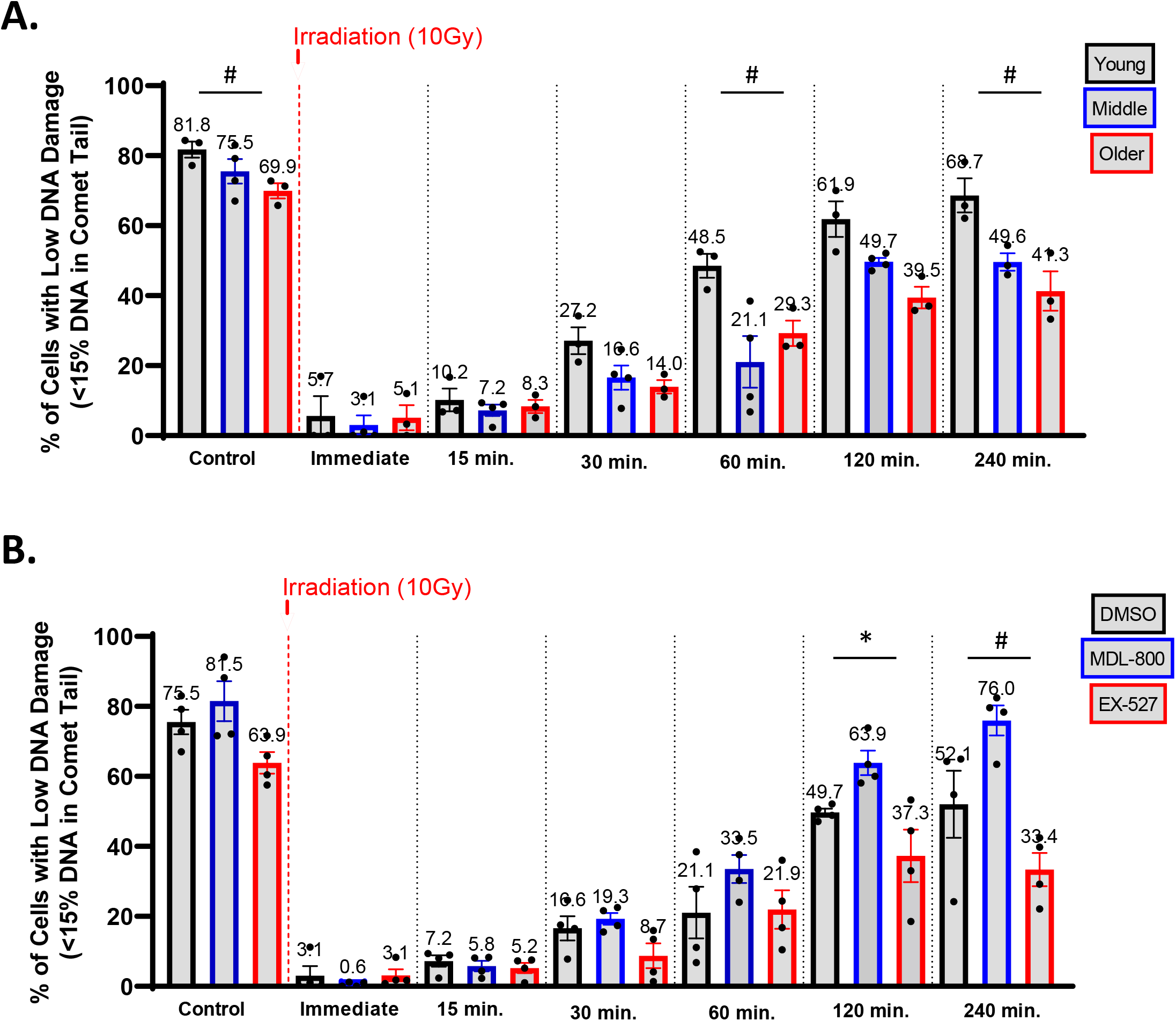
Efficacy of chondrocytes to restore low levels of DNA damage after acute damage. (A) Primary human chondrocytes from young, middle, and older donors prepared as described in Figure 1. The percentage of cells with low levels of DNA damage (<15% of DNA in comet tail) were plotted and the mean of all donors per age group is shown (mean + SEM). Repair time, age, and their interaction were significant sources of variation (2-way repeated measures ANOVA). Significant differences between groups at each time point (Tukey’s multiple comparisons test, p<0.05) are denoted by symbols: (*) = young vs. middle, (#) = young vs. old, (&) = middle vs. old. (B) Chondrocytes from middle-aged donors were pre-treated with 20 μM MDL-800, 10 μM EX-527, or vehicle (DMSO) for 2 hours before trypsinization, gel encapsulation, and irradiation. The percentage of cells with low levels of DNA damage (<15% of DNA in comet tails). Statistics similar to A, but with significant differences denoted by symbols: (*) = DMSO vs. MDL, (#) = MDL vs. EX, (&) = DMSO vs. EX).

**Figure 6:**
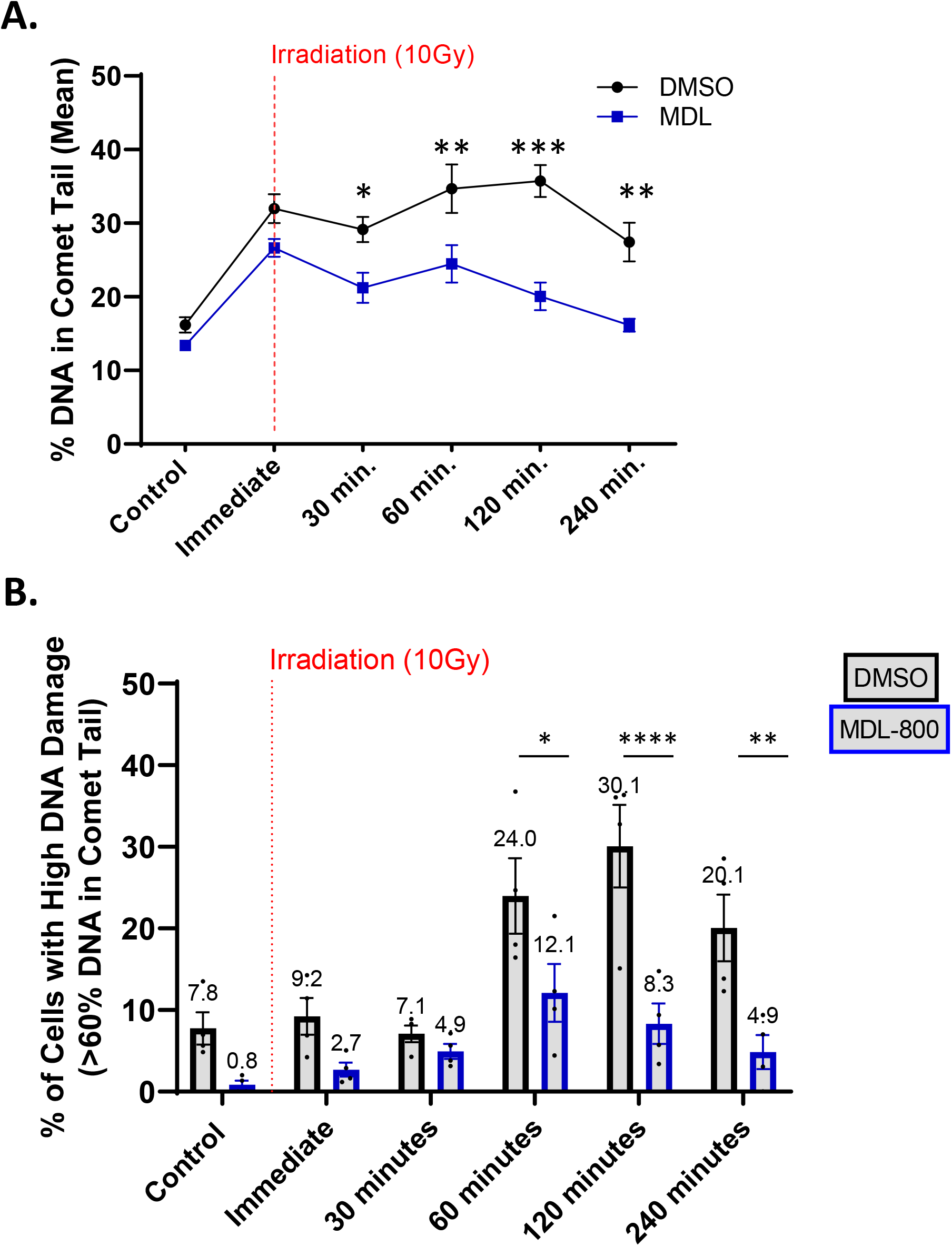
Effect of SIRT6 activation and inhibition on older chondrocyte repair efficiency. Chondrocytes from older donors (n=4, >70 years) were pre-treated with 20 μM MDL-800, 10 μM EX-527, or vehicle (DMSO) for 2 hours before trypsinization, gel encapsulation, and irradiation. Treatment continued during the repair phase. (A) The percentage of DNA in comet tails for all cells were averaged for each condition, and the mean of all donors per age group is shown (mean + SEM). Repair time, treatment, and their interaction were significant sources of variation (2-way repeated measures ANOVA). Significant differences between groups at each time point (Tukey’s multiple comparisons test, p<0.05) are denoted by symbols: (*) = DMSO vs. MDL. (B) The percentage of cells with high levels of DNA damage (>60% of DNA in comet tails). Statistics as in A.

### Statistical Analysis

Comet data were analyzed and plotted using GraphPad Prism 9. Statistical analysis was performed using paired t-test, two-way ANOVA, or two-way repeated measures ANOVA. Multiple comparison test used either Sidak’s (two treatment groups) or Tukey’s (three treatment groups) within each time point.

## Abbreviations

OA: osteoarthritis
SIRT6: Sirtuin 6
DNA: deoxyribonucleic acid
DSB: double strand breaks
SSB: single strand breaks
IR: irradiation

## Author Contributions

Conception and experimental design of this study was completed by MEC and BOD, with input from JAC. Data collection and analysis was done by MEC, JS, HLB, KRN, and BOD. Study materials were provided by SC and RFL. MEC drafted the manuscript and BOD and RFL provided critical revision and edits. All authors have approved of the final article provided herein.

## Acknowledgements

We appreciate the work of Dr. Richard Loeser’s laboratory members for help in isolating human cartilage. We acknowledge the Gift of Hope Tissue and Organ Donor Bank, the families of tissue donors, and Mrs. Arnavaz Hakimiyan for tissue procurement. Procurement of human tissue supported in part by the Rush University Klaus Kuettner Endowed Chair for Research on Osteoarthritis (SC). Funding sources had no role in the study or in the decision to publish.

